# tACS increases alpha burst duration in a frequency- and montage-specific manner

**DOI:** 10.64898/2026.08.31.747228

**Authors:** Julio Rodriguez-Larios, Anna Ragone, Satyam Chauhan, Koji Koizumi, Caroline Di Bernardi Luft

## Abstract

Transcranial alternating current stimulation (tACS) can experimentally manipulate neural oscillations in humans non-invasively. Previous studies consistently suggest that ∼10 Hz tACS in posterior regions leads to a general increase in alpha (7 – 15 Hz) oscillations. However, the specificity of this effect regarding stimulation frequency and location as well as its consequences on cognition are still unclear. We therefore aimed to investigate whether the putative effects of tACS on neural oscillations and cognition are specific to posterior stimulation in the alpha frequency range. We tested these effects using a 2 x 2 design (posterior or fronto-parietal stimulation, at 10 Hz or 6 Hz) plus a no-stimulation control group, during a sustained attention task.158 healthy participants were randomly allocated to control group (no stimulation) or one of the four active groups. All participants underwent an Electroencephalography recording and completed the Amsterdam Resting State Questionnaire before and after stimulation. Our results show that, relative to the control group, alpha power increases significantly after stimulation for the group receiving posterior 10 Hz stimulation and that this was explained by an increase in alpha oscillatory bursts duration rather than their amplitude. On the other hand, no significant differences were found between active and controls groups in behavioural performance or resting state cognition. Together, our findings demonstrate that tACS increases the duration of alpha oscillatory bursts in a montage- and frequency-specific manner and that this modulation is not accompanied by significant changes in sustained attention during stimulation or resting state cognition after stimulation .

## Introduction

Neural oscillations are rhythmic patterns of electrical activity coming from the brain (Buzsáki & Draguhn, 2004). These rhythms are thought to be generated by summed dendritic postsynaptic potentials (Buzsáki et al., 2012), although the exact origin might differ depending on the involved network (Halgren et al., 2019; Sherman et al., 2016). Several studies in humans have shown that neural oscillations play a key role in healthy cognitive function (Klimesch, 1999) and are affected in clinical populations with cognitive deficits (Babiloni et al., 2025; Uhlhaas & Singer, 2006). Because the relationship between neural oscillations and cognition is largely based on correlational evidence, neuromodulation techniques have attracted considerable interest as a tool to manipulate oscillatory activity and this way test its putative causal role in cognition (Riddle & Frohlich, 2021).

Transcranial alternating current stimulation (t ACS) is a non-invasive neuromodulation technique consisting of the delivery of a rhythmic electrical current to the brain through the scalp (Wischnewski et al., 2022), which has been shown to be able to modulate brain activity online (during stimulation) (Violante et al., 2017) and offline (after stimulation) (Grover et al., 2023; Veniero et al., 2015). Several studies demonstrated that tACS can specifically modulate neural oscillations (Wischnewski et al., 2023). The most consistent finding is the increase of alpha oscillatory power (7 – 15 Hz) after ∼10 Hz tACS is delivered to posterior areas for approximately 20 minutes (De Koninck et al., 2021; Helfrich et al., 2014; Neuling et al., 2017; Vossen et al., 2015; Zaehle et al., 2010a). Although the mechanisms behind this effect are still debated, the dominant account combines entrainment during stimulation with plasticity after it. Rhythmic electrical stimulation is thought to bias the timing of neural spiking towards the phase of the applied current, but only when the stimulation frequency is sufficiently close to that of an ongoing endogenous rhythm (W. A. Huang et al., 2021; Krause et al., 2019, 2022). Repeated coincidence between stimulated and endogenous activity would then strengthen the synaptic connections sustaining that rhythm via spike-timing-dependent plasticity (STDP), producing the after-effects observed once stimulation has ceased (Agboada et al., 2025; Wischnewski et al., 2023). This account makes two predictions. First, after-effects should depend on the match between stimulation frequency and the endogenous rhythm. Second, they should depend on the presence of that rhythm in the stimulated tissue, and therefore on where the current is delivered.

The frequency and spatial specificity of the reported tACS effects on alpha oscillations are largely unexplored. Although some computational work suggest that stimulation at the endogenous frequency (or slightly lower) would result in greater increase in alpha power after stimulation (Vossen et al., 2015; Zaehle et al., 2010b), this has not been systematically tested. In fact, alpha power increases after tACS have been reported for both individualised alpha frequency (Clancy et al., 2018; Kasten et al., 2016) and for a predefined frequency within the alpha range (e.g.10 Hz) (Clayton et al., 2019). In the same way, no previous study investigated whether tACS can affect alpha oscillations when delivered at a different location. Since different alpha rhythms have been found throughout the cortex (Haegens et al., 2011; Rodriguez-Larios et al., 2022; Sokoliuk et al., 2019; van der Vinne et al., 2017), similar alpha after-effects could be expected when tACS is delivered to central and frontal areas.

A second open question concerns the nature of the alpha increase itself. Alpha power is typically quantified over recordings lasting several minutes, which obscures the fact that alpha activity is not sustained but occurs in transient bursts (Ossadtchi et al., 2017; Vidaurre et al., 2016, 2018). Because of this, a given increase in average power can arise in several ways: bursts may become stronger, longer, or more frequent. These possibilities are not equivalent, since they plausibly reflect different physiological processes and may carry different functional consequences (Bonaiuto et al., 2021; Jones et al., 2009). Characterising tACS after-effects at the level of burst dynamics therefore offers a more mechanistically informative description than average power alone. To our knowledge, this has not been attempted for tACS after-effects.

The consequences of alpha tACS on cognition are still controversial. An important number of studies reporting alpha power changes after tACS include a sustained attention task during stimulation (De Koninck et al., 2021; Helfrich et al., 2014; Neuling et al., 2017; Vossen et al., 2015; Zaehle et al., 2010a). However, only some studies report effects on performance. Specifically, it has been suggested that posterior alpha tACS can stabilise performance in different attention tasks but these results are not fully consistent (Clayton et al., 2018, 2019; Klink et al., 2020). On the other hand, the potential effects of tACS on subjective cognitive state after stimulation have received considerably less attention. Assessing resting-state cognition after stimulation may provide a complementary measure of whether changes in neural oscillations are accompanied by changes in how participants experience their own cognitive state.

In this study, we investigated whether the putative effects of tACS on neural oscillations and cognition are specific to posterior stimulation in the alpha frequency range. For this purpose, we delivered tACS during a sustained attention task (Psychomotor Vigilance Task; PVT) in four active groups (posterior or fronto-parietal stimulation at 10 Hz or 6 Hz) and a control group (sham stimulation). We recorded Electroencephalography (EEG) and characterised changes in alpha oscillations via a recently developed algorithm (Rodriguez-Larios et al., 2024; Rodriguez-Larios C Haegens, 2023). In addition, we quantified resting state cognition before and after stimulation through the Amsterdam Resting State Questionnaire (ARSQ). We specifically assess which active group shows a significant change in alpha oscillations, behaviour and/or resting state cognition relative to the control group.

## Methods

### Participants

A total of 158 healthy adults (93 females; mean age = 23.3 years, SD = 5.3) participated in the study. Participants were recruited from the Brunel University of London campus, the surrounding community, and via the University’s SONA participant recruitment system. Exclusion criteria included a history of neurological disorders (including epilepsy), the presence of implanted medical devices, current use of psychotropic medication, or any other contraindication to transcranial electrical stimulation. All participants had normal or corrected-to-normal vision and provided written informed consent prior to participation. Participants received either £20 or course credit for their participation. The study was approved by the College of Health, Medicine and Life Sciences Research Ethics Committee at Brunel University of London (Reference No. 47845) and was conducted in accordance with the Declaration of Helsinki.

Participants were randomly allocated to one of five experimental groups: control (no stimulation, *n* = 30), fronto-parietal alpha stimulation (*n* = 30), fronto-parietal theta stimulation (*n* = 28), posterior alpha stimulation (*n* = 33), and posterior theta stimulation (*n* = 33). Note that sample size was different for the analysis of each of the 3 dependent variables (questionnaire, performance and EEG) as some subjects had to be excluded due to some technical problems during data acquisition:

- The ARSQ was analysed in 152 subjects (30 control, 29 fronto-parietal alpha stimulation, 27 fronto-parietal theta stimulation, 33 posterior alpha stimulation, and 33 posterior theta stimulation).
- PVT performance was analysed in 146 subjects (29 control, 30 fronto-parietal alpha stimulation, 28 fronto-parietal theta stimulation, 29 posterior alpha stimulation, and 30 posterior theta stimulation).
- EEG was analysed in 144 subjects (30 control, 29 fronto-parietal alpha stimulation, 27 fronto-parietal theta stimulation, 30 posterior alpha stimulation, and 28 posterior theta stimulation).

### Design, procedure and task

The experiment consisted of a pre-stimulation resting-state EEG recording (5 minutes), a 20-minutes tACS session during PVT performance, and a post-stimulation resting-state EEG recording (5 minutes) (see **Figure 1A)**. Resting-state EEG was recorded while participants remained seated with their eyes open and fixated on a centrally presented cross. Following each resting-state recording, participants completed the ARSQ (Diaz et al., 2013).

**Figure 1.**
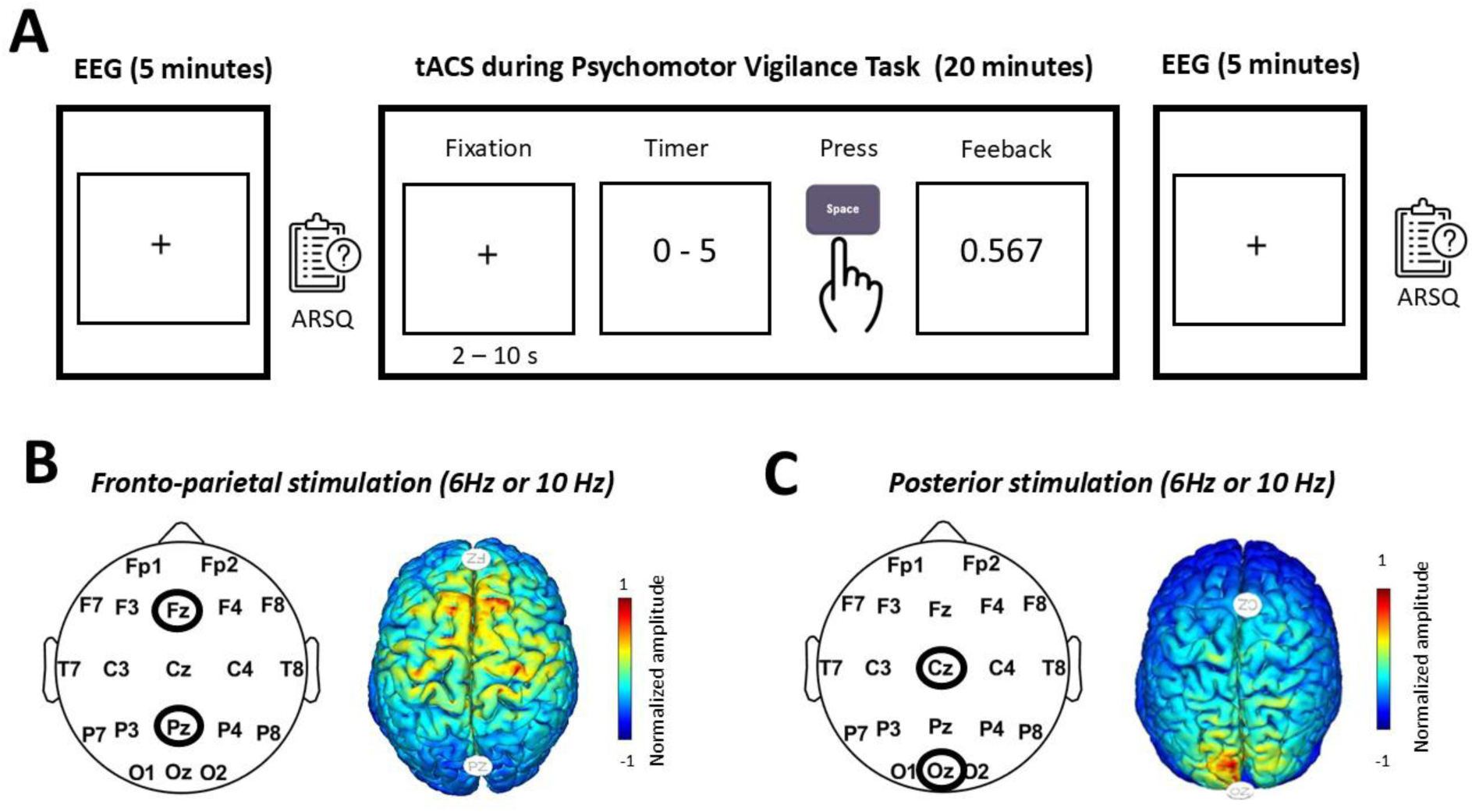
Experimental design and transcranial alternating current stimulation (tACS) protocols. (A) Experimental procedure. Participants completed a 5-minute resting-state EEG recording before and after tACS while fixating on a central cross. Immediately following each resting-state recording, participants completed the ARSQ. During the 20-minute tACS session, participants performed PVT. (B) Fronto-parietal stimulation protocol. The left panel shows the EEG electrode layout, with the stimulation electrodes positioned over Fz and Pz (circled). The right panel shows the corresponding modelled electric field distribution, displayed as normalised electric field amplitude across the cortical surface. (C) Posterior stimulation protocol. The left panel shows the EEG electrode layout, with the stimulation electrodes positioned over Cz and Oz (circled). The right panel shows the corresponding modelled electric field distribution, displayed as normalised electric field amplitude across the cortical surface. For both stimulation montages, separate groups received either 10 Hz (alpha) or 6 Hz (theta) tACS.

During the stimulation session, participants performed a 20-min Psychomotor Vigilance Task (PVT)(Lim & Dinges, 2008) while receiving tACS. The PVT is a sustained attention task in which participants continuously monitor a fixation point for the appearance of a millisecond counter and are instructed to respond as quickly as possible upon its onset while avoiding anticipatory responses. The inter-trial interval was between 2 and 10 seconds, which resulted in a total of 160 trials approximately. The task was implemented in MATLAB R2024a (MathWorks, Natick, MA, USA) using Psychtoolbox and was presented on a Windows 10 computer.

### EEG acquisition and tACS delivery

Simultaneous EEG and tACS were delivered using a Neuroelectrics Starstim system (Neuroelectrics, Barcelona, Spain). EEG was recorded from 20 Ag/AgCl electrodes positioned according to the international 10–20 system, with the reference electrode placed on the right side of the neck. tACS was delivered through two stimulation montages: Cz–Oz (posterior stimulation) and Fz–Pz (fronto-parietal stimulation) (see **Figure 1B-C**). Depending on the experimental group, stimulation was applied at either 6 Hz (theta) or 10 Hz (alpha), resulting in four active stimulation protocols (posterior theta, posterior alpha, fronto-parietal theta, and fronto-parietal alpha). Participants in the control group underwent the same experimental procedure but did not receive tACS.

Prior to stimulation, a comfort-guided intensity titration procedure was used to determine each participant’s maximum comfortable stimulation intensity. Stimulation commenced at 0.5 mA and was increased in 0.5 mA increments to a maximum of 2.0 mA (0-to-peak; 4 mA peak-to-peak). Following each increment, participants rated the comfort of the stimulation, and the titration was terminated when the stimulation became uncomfortable. The stimulation intensity used during the experimental session was the highest amplitude that each participant judged to be comfortable.

### EEG preprocessing

The data were band-pass filtered between 1 and 30 Hz using zero-phase finite impulse response filters. Flat channels (>60 s of flat signal) and noisy channels identified by low correlation with a robust channel estimate were automatically detected and removed. Transient high-amplitude artefacts were attenuated using the Artifact Subspace Reconstruction (ASR) algorithm (Plechawska-Wojcik et al., 2019) after which removed channels were reconstructed by spherical spline interpolation. Independent component analysis (ICA) was then performed using the RUNICA algorithm with dimensionality adjusted according to the data rank. Independent components were automatically classified using ICLabel (Pion-Tonachini et al., 2019), and components classified as ocular, muscle, cardiac, or channel-noise artefacts with a classification probability greater than 80% were removed.

Power spectral density (PSD) was estimated for each resting-state EEG recording using Welch’s method with 1 second windows, 50% overlap, and a frequency resolution of 0.1 Hz (1–30 Hz). Alpha oscillatory activity was quantified separately for each electrode by identifying the largest spectral peak within the alpha band (7–15 Hz). Individual alpha frequency (IAF) was defined as the frequency of this peak, whereas alpha power was quantified as the prominence of the spectral peak, rather than its absolute amplitude. Peak prominence was defined as the vertical distance between the peak and its lowest contour, providing a measure of oscillatory strength that is less influenced by the underlying aperiodic spectral background (Donoghue et al., 2021).

Alpha oscillatory activity was further characterised with a recently developed burst detection algorithm (Rodriguez-Larios et al., 2024; Rodriguez-Larios & Haegens, 2023). In short, EEG data were transformed into the time-frequency domain using 6-cycle Morlet wavelets (1-Hz frequency resolution, 1–30 Hz). Oscillatory bursts were defined as periods during which the amplitude at a given frequency exceeded the estimated aperiodic (1/f) background for at least one oscillatory cycle. The aperiodic component was estimated for each electrode by fitting a linear function to the power spectrum in log-log space. Only bursts whose peak spectral frequency fell within the alpha band (7–15 Hz) were retained for further analysis. The algorithm quantified burst amplitude, duration, number and coverage.

### Questionnaires

The Amsterdam Resting State Questionnaire was employed to measure resting state cognition (Diaz et al., 2013). The ARSQ is a validated self-report questionnaire that measures the content and quality of resting-state cognition across multiple domains, including Discontinuity of Mind, Theory of Mind, Self, Planning, Sleepiness, Comfort and Somatic Awareness. Participants completed the questionnaire immediately following each resting-state EEG recording to characterise their mental experiences during the preceding rest period.

### Statistical analyses

Linear mixed models and paired-samples t-tests (MATLAB R2024b implementation) were used to assess the effect of tACS on behaviour, alpha oscillations and resting state cognition. The linear mixed models were specifically aimed to assess whether each stimulation group differed from the control group in any of the dependent variables. The statistical approach for each dependent variable is described below.

#### Behavioural data

Trial-level reaction times were analysed rather than participant averages to account for within-subject variability and to maximise statistical power. Trials corresponding to false alarms or responses exceeding 2 seconds were excluded from the analysis. The remaining reaction times were modelled with fixed effects of stimulation group (control, fronto-parietal alpha, fronto-parietal theta, posterior theta, and posterior alpha), trial number, and their interaction. The control group was specified as the reference category, such that the coefficients for each active stimulation group represented differences relative to the control group. Random intercepts and random slopes for trial were included for each participant to account for individual differences in baseline reaction time and the rate of change in performance across the task. Statistical significance was set at *p* < .05.

#### EEG alpha oscillations

Separate models were fitted for each electrode. Fixed effects included stimulation group (control, fronto-parietal alpha, fronto-parietal theta, posterior theta, and posterior alpha), time (pre- vs. post-stimulation), and their interaction. The control group and pre-stimulation measurements were specified as the reference categories, such that each Group × Time interaction quantified whether the pre- to post-stimulation change in alpha power differed between an active stimulation protocol and the control group. Subject was included as a random intercept to account for repeated measurements within participants

For stimulation protocols that produced significant changes in alpha power, two exploratory analyses were conducted to further characterise the observed effects. First, to investigate whether changes in alpha power were influenced by stimulation parameters or individual alpha frequency (IAF), Pearson correlation analyses were performed. Alpha power change was calculated as the post-minus pre-stimulation difference and averaged across electrodes showing a significant Group × Time interaction. Three variables were examined: (1) stimulation amplitude (mA), (2) the absolute mismatch between each participant’s IAF and the stimulation frequency (i.e., |IAF − 10 Hz|), and (3) mean IAF (averaged across the significant electrodes). Pearson correlation coefficients were computed between each predictor and the mean change in alpha power. Second, to characterise the underlying oscillatory dynamics associated with the observed alpha power changes (Rodriguez-Larios et al., 2024; Rodriguez-Larios & Haegens, 2023), four burst metrics were quantified: burst amplitude, burst duration, burst number, and burst coverage. For each metric, values were averaged across the alpha frequency range (7–15 Hz) and compared between pre- and post-stimulation sessions using paired-samples *t*-tests at each electrode. False discovery rate (FDR) correction was applied across electrodes to control for multiple comparisons.

#### Amsterdam Resting State Questionnaire

Separate models were fitted for each ARSQ dimension, with questionnaire score entered as the dependent variable. Fixed effects included stimulation group (control, fronto-parietal alpha, fronto-parietal theta, posterior theta, and posterior alpha), time (pre- vs. post-stimulation), and their interaction. The control group and pre-stimulation measurements were specified as the reference categories, such that each Group × Time interaction quantified whether the pre- to post-stimulation change in questionnaire scores differed between an active stimulation protocol and the control group. Subject was included as a random intercept to account for repeated measurements within participants. Model coefficients, *t*-statistics, and *p*-values were extracted separately for each ARSQ dimension.

## Results

### Psychomotor Vigilance Task

As shown in **Figure 2**, reaction times increased progressively over the course of the task, consistent with the expected vigilance decrement (Lim & Dinges, 2008). Across all participants and trials, the mean reaction time was 0.4 s (*SD* = 0.09 s). A linear mixed-effects model revealed a significant effect of Time, *β* = 0.00053, *SE* = 0.00015, *t*(22,272) = 3.13, *p* < .001, confirming a progressive slowing of responses over the course of the task. There was no significant main effect of Group (all *ps* > .05), and none of the Group × Trial interactions were significant (all *ps* > .05), indicating that neither overall reaction times nor the rate of vigilance decline differed significantly between any of the active stimulation protocols and the control group. The full model outputs, including fixed-effect coefficients, *t*-statistics, and *p*-values, are provided in **Supplementary Table 1**.

**Figure 2.**
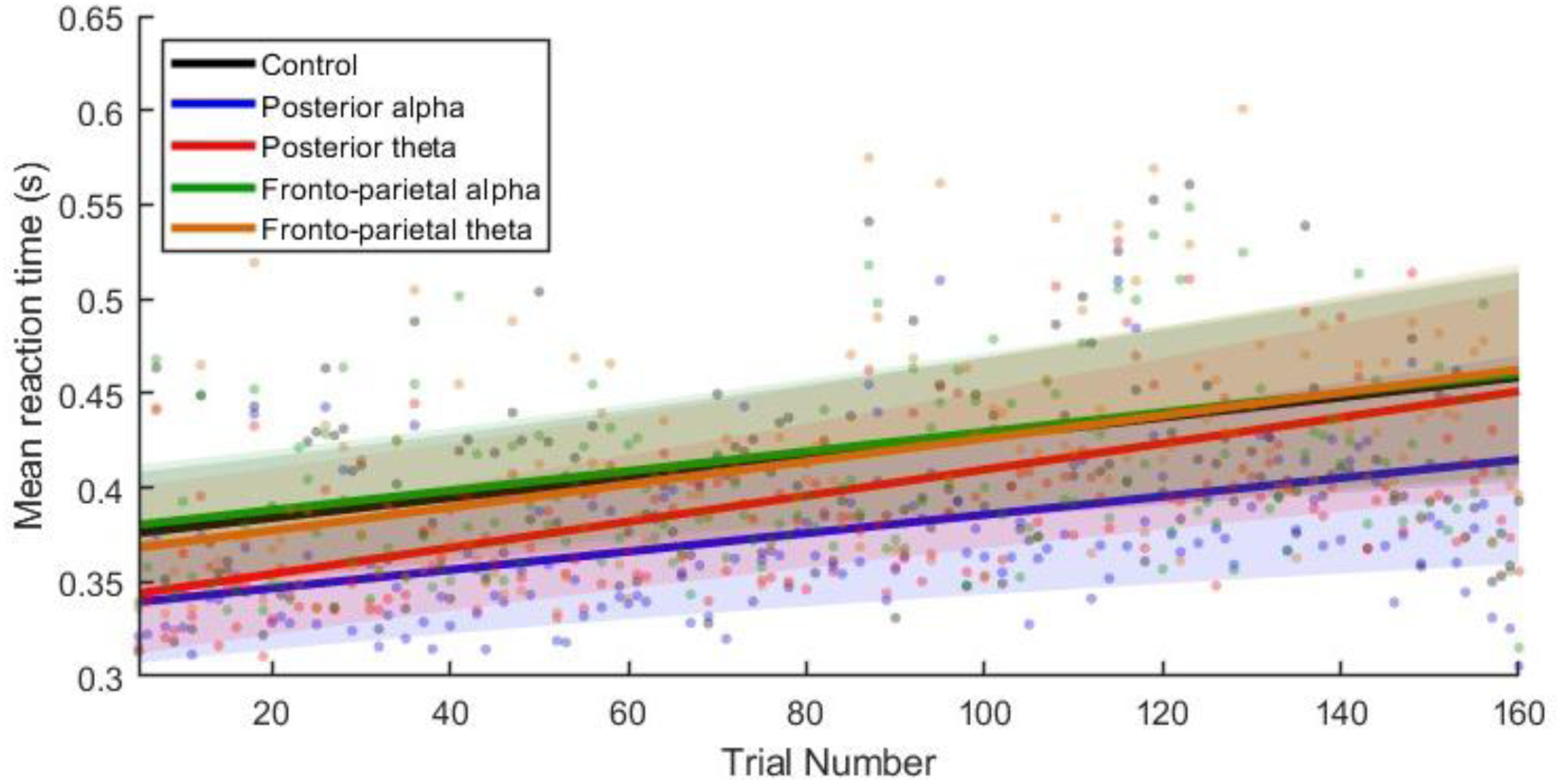
Behavioural performance during the Psychomotor Vigilance Task. Mean reaction time for each trial and group is shown as scatter points, and the fitted trajectories from the linear mixed-effects model are shown as solid lines. Shaded regions represent the 95% confidence intervals of the model predictions. The control group is shown in black, the posterior alpha stimulation group in blue, the posterior theta stimulation group in red, the fronto-parietal alpha stimulation group in green, and the fronto-parietal theta stimulation group in orange. Although reaction times increased over the course of the task across all groups, no significant differences in reaction time or rate of change over time were observed between any active stimulation protocol and the control group.

### Resting state cognition

Linear mixed-effects models were fitted separately for each ARSQ dimension to examine Time x Protocol interactions. No significant Time × Protocol interactions were observed for any ARSQ dimension (all *ps* > .05), indicating that changes in self-reported resting-state cognition from pre- to post-stimulation did not differ between any of the active stimulation protocols and the control group. The full model outputs, including fixed-effect coefficients, *t*-statistics, and *p*-values for each ARSQ dimension, are provided in the Supplementary Materials (see **Supplementary Tables 2-8**).

### Alpha power analyses

A significant Group × Time interaction (*p_FDR_* < .05) was only identified for the posterior alpha stimulation protocol (i.e., Oz–Cz stimulation at 10 Hz), indicating that the pre- to post-stimulation change in alpha power differed significantly from that of the control group. No significant Group × Time interactions were found for the fronto-parietal alpha, fronto-parietal theta, or posterior thetastimulation protocols. Note that although some differences were found for the fronto-parietal theta protocol, these did not survive FDR correction (see **Supplementary Tables 9-29** for mixed-model results of each electrode). The topographical distribution of the interaction effects and the corresponding pre- and post-stimulation power spectra are shown in **Figure 3A-H**.

**Figure 3.**
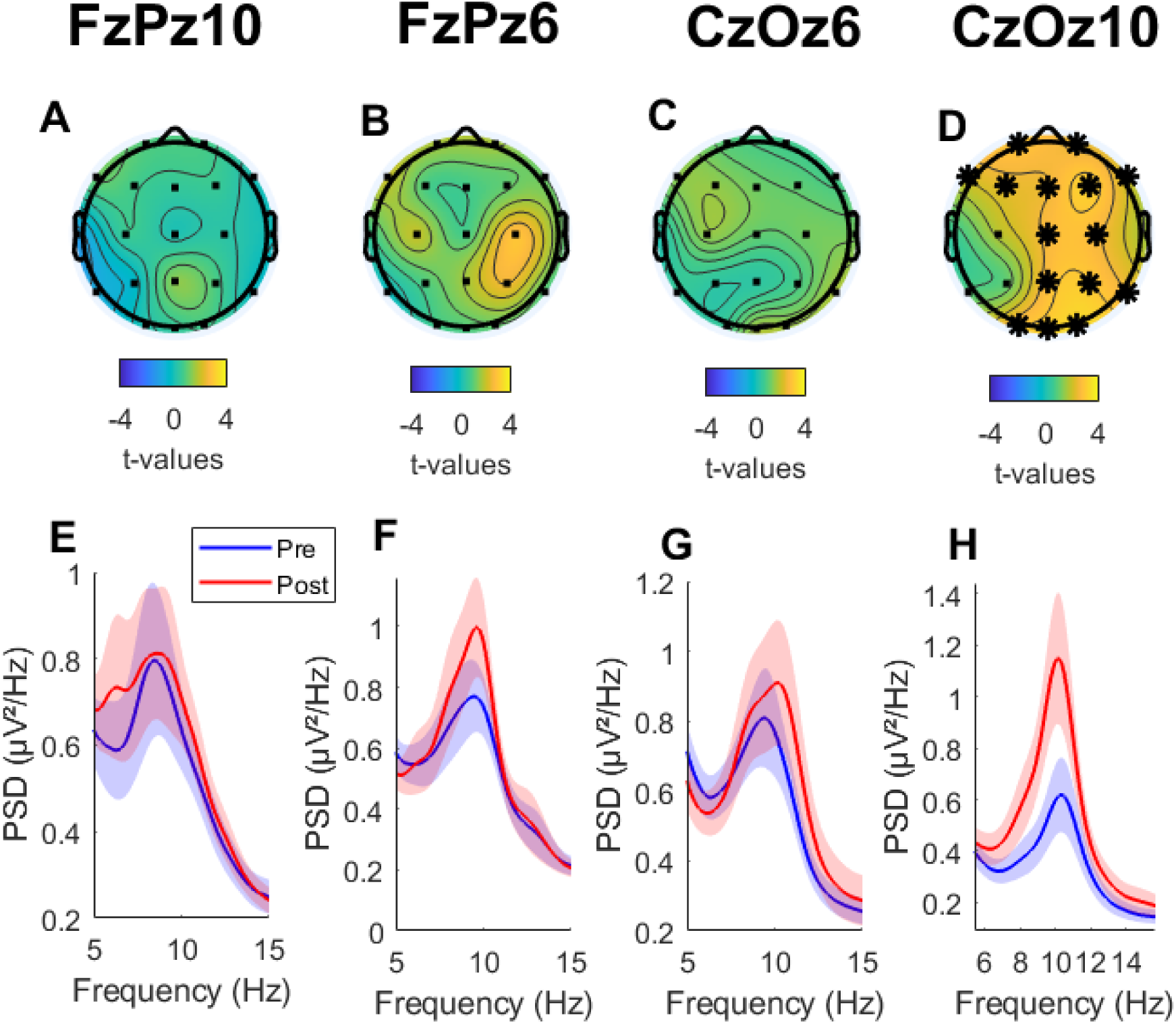
Group × Time interaction effects on alpha power following transcranial alternating current stimulation (tACS). **(A–D)** Topographical maps showing the *t*-values of the Group × Time interaction from the linear mixed-effects models for each active stimulation protocol relative to the control group: fronto-parietal alpha (FzPz10), fronto-parietal theta (FzPz6), posterior theta (CzOz6), and posterior alpha (CzOz10). Asterisks indicate electrodes showing a significant interaction after false discovery rate (FDR) correction (*p* < .05). **(E–H)** Mean power spectral density before (Pre, blue) and after (Post, red) stimulation for each protocol. Shaded regions represent ±1 standard error of the mean. Spectra were averaged across electrodes showing significant interaction effects for the Posterior alpha protocol and across all electrodes for the remaining stimulation protocols.

On the other hand, exploratory analyses revealed no significant associations between the change in alpha power and any of the stimulation-related variables (**Figure 4**). Specifically, alpha power change was not significantly correlated with stimulation amplitude (*r* = −.23, *p* = .213), the absolute mismatch between IAF and the 10 Hz stimulation frequency (*r* = −.29, *p* = .126), or participants’ IAF (*r* = −.12, *p* = .521). These findings suggest that the increase in alpha power following posterior alpha stimulation was not explained by individual differences in stimulation intensity or baseline alpha frequency.

**Figure 4.**
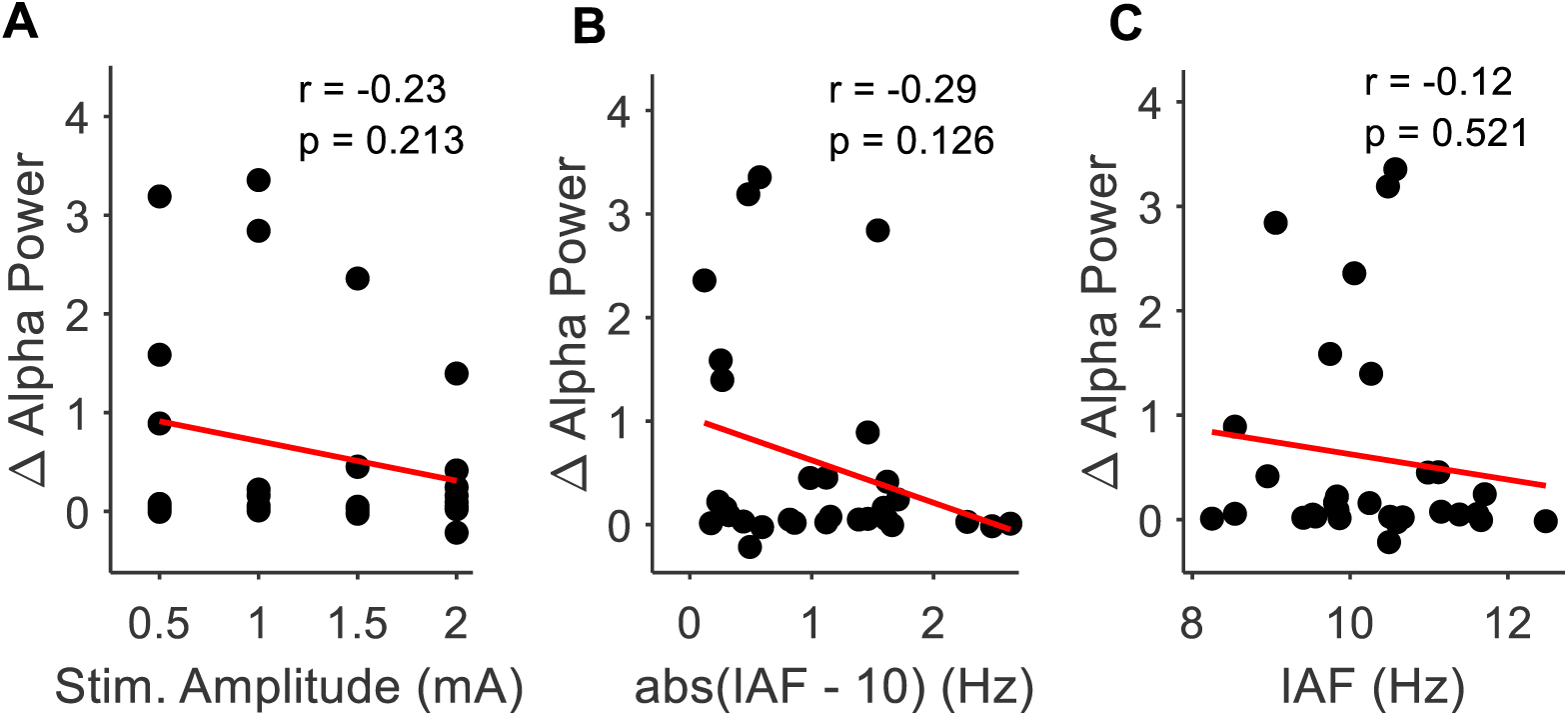
Exploratory correlations between the change in alpha power following posterior alpha tACS and stimulation-related variables. Alpha power change(post − pre) was averaged across electrodes showing a significant Group × Time interaction. Scatterplots depict the relationship between alpha power change and **(A)** stimulation amplitude, **(B)** the absolute mismatch between individual alpha frequency (IAF) and the 10 Hz stimulation frequency (|IAF− 10|), and **(C)** IAF. Red lines indicate the least-squares regression fit. Pearson correlation coefficients (*r*) and corresponding *p*-values are shown in each panel. No significant correlations were observed.

### Oscillatory bursts analyses

To characterise the oscillatory dynamics underlying the observed increase in alpha power in the posterior alpha stimulation group, four alpha burst metrics were examined: burst amplitude, burst duration, burst number, and burst coverage (**Figure 5**). No significant changes were observed in burst amplitude or burst number following stimulation (**Figures 5A–B** and **Figures 5E–F**, respectively). In contrast, alpha burst duration increased significantly from pre- to post-stimulation (**Figures 5C–D**). Likewise, burst coverage (the proportion of time occupied by alpha bursts) increased significantly after stimulation (**Figures 5G–H**). Together, these findings indicate that the observed increase in alpha power following posterior alphatACS was primarily driven by longer-lasting and more prevalent alpha bursts, rather than increases in burst amplitude or the number of bursts.

**Figure 5.**
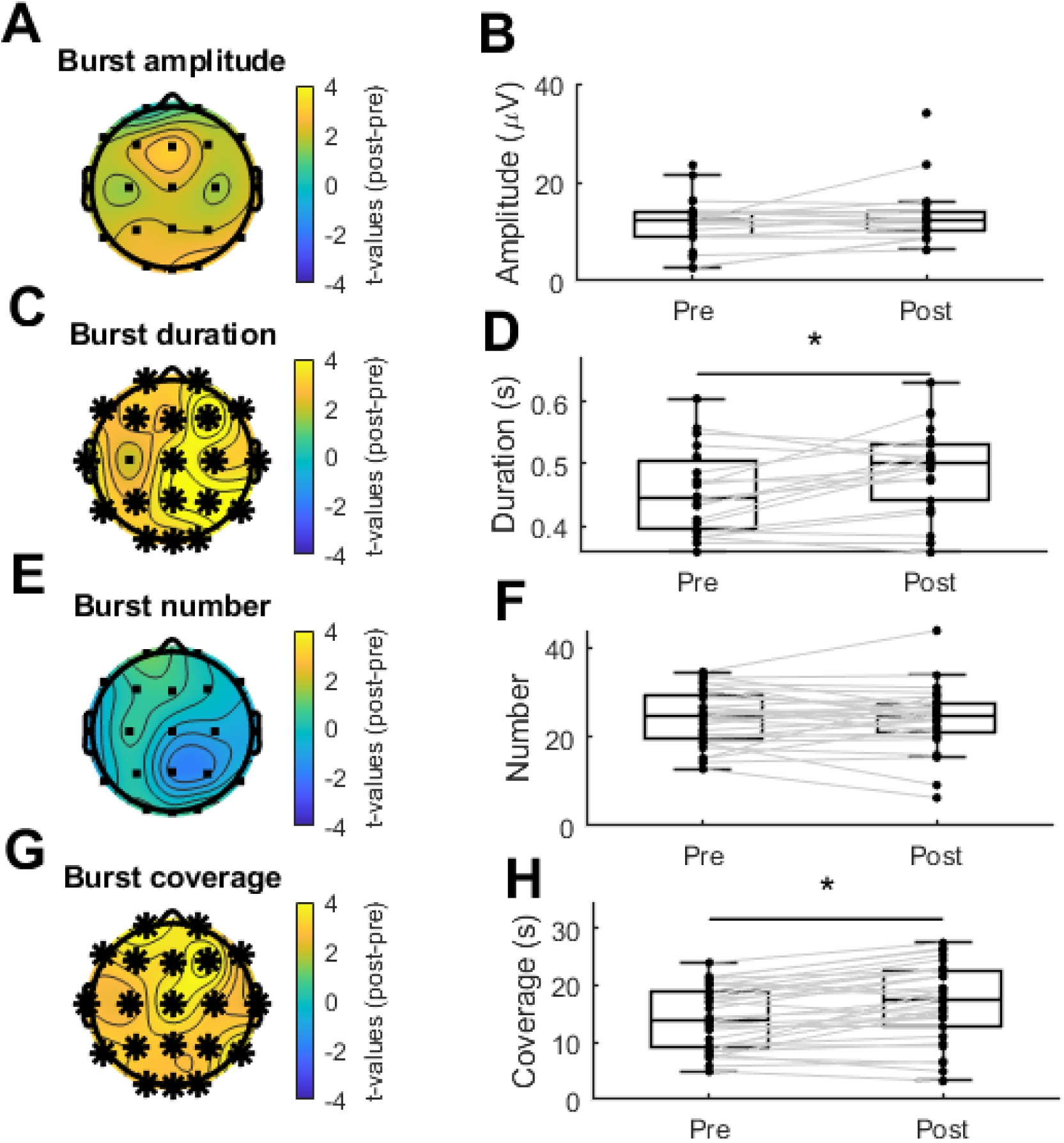
Burst dynamics following posterior alpha tACS. Changes in alpha burst characteristics following posterior alpha stimulation (Oz–Cz, 10 Hz). Left panels show topographical distributions of paired-samples *t*-values (post − pre) for **(A)** burst amplitude, **(C)** burst duration, **(E)** burst number, and **(G)** burst coverage. Electrodes showing significant pre- to post-stimulation differences after false discovery rate (FDR) correction are marked with black asterisks. Right panels show participant-level paired boxplots averaged across significant electrodes for **(B)** burst amplitude, **(D)** burst duration, **(F)** burst number, and **(H)** burst coverage. Each dot represents an individual participant, with grey lines connecting pre- and post-stimulation measurements. Significant increases were observed for burst duration and burst coverage, whereas burst amplitude and burst number did not differ significantly between sessions.

## Discussion

In this study, we assessed the effects of different tACS protocols on behaviour, resting state cognition and alpha oscillations. Specifically, we tested the effects of posterior and fronto-parietal stimulation at alpha (10 Hz) and theta (6 Hz) frequencies in four different groups by comparing them to a control group that did not receive stimulation. None of the groups differed significantly from the control group in terms of behaviour (i.e. performance in vigilance task) or resting state cognition (i.e. Amsterdam Resting State Questionnaire. Regarding neural effects, alpha power was only significantly increased after stimulation in theposterior alphastimulation group while no significant effects for fronto-parietal alpha, fronto-parietal theta nor posterior theta groups were found. The alpha power increase in the posterior alpha group did not depend on stimulation amplitude nor individual alpha frequency. In addition, oscillatory bursts analyses revealed that the alpha power increase in the posterior alpha group was due to an increase in oscillatory burst duration and coverage rather than a change in oscillation amplitude.

In line with previous literature, we found that posterior 10 Hz tACS increases EEG alpha power after stimulation (De Koninck et al., 2021; Helfrich et al., 2014; Neuling et al., 2017; Vossen et al., 2015; Zaehle et al., 2010a). In addition, our results revealed that this effect has some degree of spatial and frequency specificity by showing no significant effects with fronto-parietal stimulation (at 6 Hz or 10 Hz) nor posterior stimulation at 6 Hz. The spatial and frequency specificity of tACS on alpha oscillations can be interpreted in the light of the entrainment theory (Riddle & Frohlich, 2021). According to this theory, rhythmic electric stimulation will only entrain neural spiking activity when delivered at the frequency of the endogenous neural rhythm (W. A. Huang et al., 2021; Krause et al., 2019, 2022), which is thought to drive tACS after-effects via STDP(Agboada et al., 2025). Since posterior areas are dominated by alpha rhythms between 7 and 15 Hz (Haegens et al., 2014; Rodriguez-Larios et al., 2022; Sokoliuk et al., 2019), alpha entrainment after 10 Hz stimulation would be expected. Nonetheless, it is important to highlight that our reported alpha effects did not depend on individual alpha frequency (see **Figure 4 B-C**), suggesting that entrainment of posterior alpha rhythms through tACS could occur in a wide range of frequencies within the alpha band. Therefore, future studies in which stimulation frequency is parametrically modulated are needed to estimate the exact posterior alpha tACS stimulation frequency limits.

Burst analysis have not been previously performed to assess tACS after-effects on neural oscillations. By applying a recently developed algorithm (Rodriguez-Larios et al., 2024; Rodriguez-Larios & Haegens, 2023), we show that alpha power increases after posterior 10 Hz tACS is due to an increase duration and coverage of oscillatory bursts rather than an increase in burst amplitude. This dissociation may provide novel insights into the physiological mechanisms underlying alpha generation (Bastiaens et al., 2025; Vijayan & Kopell, 2012). In this context, combining controlled perturbations of neural oscillations through tACS with oscillatory burst analyses may provide a useful experimental framework for constraining and refining future computational models of alpha generation. Furthermore, distinguishing the effects of tACS on different oscillatory burst properties may have important implications for clinical applications of neuromodulation. Different neurological and psychiatric disorders may be characterised by alterations in distinct aspects of oscillatory dynamics, such as burst occurrence, duration, or amplitude, and these properties may respond differently to specific stimulation protocols. For example, we have previously shown that ADHD traits are associated with a reduced occurrence of alpha bursts, while burst amplitude remains preserved (Rodriguez-Larios et al., 2025). Based on the present findings, it would be relevant to determine whether increasing the coverage of alpha activity through tACS translates into behavioural improvements in individuals with ADHD.

The null effects of tACS on cognitive performance in this study partially contradict previous literature. Specifically, it has been shown that alpha tACS stabilises sustained attention performance in different tasks (Clayton et al., 2019). Two methodological differences may account for this discrepancy. First, although both studies assessed sustained attention, the behavioural paradigms differed substantially. Whereas Clayton et al. (2019) employed continuous visual attention tasks requiring sustained perceptual processing, the present study used the PVT, which primarily measures vigilance. Given that posterior alpha oscillations have been most consistently implicated in visual attention and perceptual processing (Pascucci et al., 2025), it is possible that alpha tACS preferentially modulates these functions rather than vigilance per se. Second, in Clayton et al. (2019) behavioural effects emerged during the final five minutes of the task after stimulation had ceased, whereas stimulation in the present study was delivered throughout the entire task. It is therefore possible that behavioural effects of posterior alpha tACS are more prominent after stimulation has ended, in parallel with the offline increase in alpha power observed in the present study.

This study has several limitations. First, online neural effects could not be assessed because of the large stimulation artefacts introduced by tACS, limiting our ability to determine the mechanisms underlying the observed after-effects. Consequently, we cannot establish whether posterior alpha tACS enhances endogenous alpha oscillations during stimulation through neural entrainment (Helfrich et al., 2014), or whether the post-stimulation increase in alpha power reflects a rebound phenomenon following stimulation (Haberbosch et al., 2019). Recent methodological developments in simultaneous EEG-tACS recordings may help address this question in future studies (Ros et al., 2025). Second, individual MRI-based electric-field modelling was not performed (Caulfield & George, 2022; Y. Huang et al., 2019). Given that anatomical variability can substantially influence current distribution and tACS efficacy (Kasten et al., 2019), incorporating individualised electric-field modelling may help explain inter-individual variability in stimulation responses. Finally, the present study does not address the influence of behavioural context on tACS after-effects. Because stimulation was delivered while participants performed a vigilance task, it remains unknown whether similar changes in alpha burst dynamics would occur during resting-state stimulation or other cognitive tasks. Identifying the behavioural contexts that maximize the effects of alpha tACS will be particularly important if this technique is to be translated into clinical applications (Agboada et al., 2025).

In conclusion, the present study demonstrates that posterior 10 Hz tACS selectively increases resting-state alpha power in a montage- and frequency-specific manner. In addition, burst analyses revealed that this effect is driven by greater coverage and duration of alpha bursts rather than increased burst amplitude. These findings provide new insight into the neural dynamics underlying tACS after-effects and highlight oscillatory burst analyses as a promising approach for investigating the mechanisms and potential clinical applications of non-invasive brain stimulation.

---

**Supplementary Table 1.**
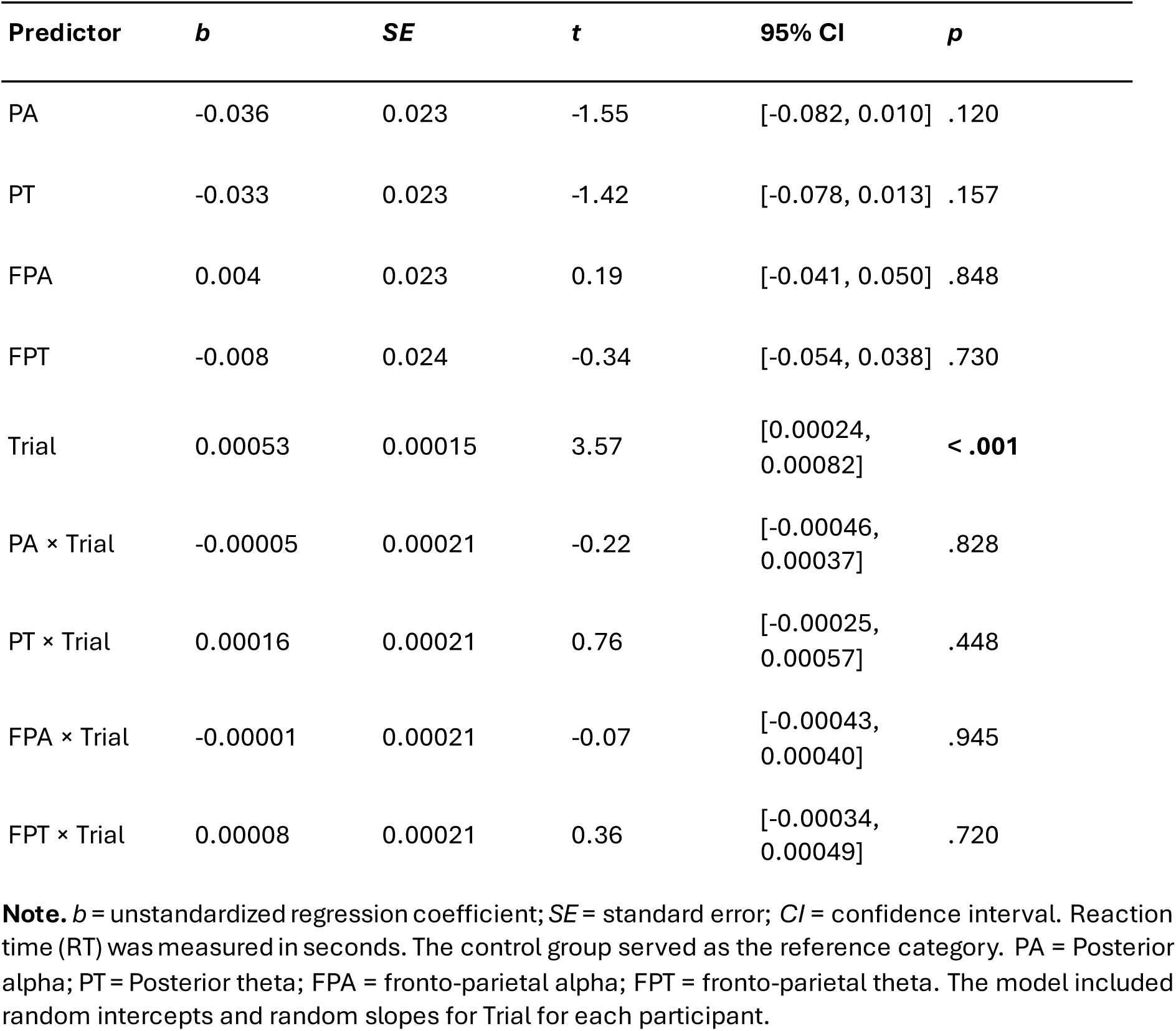
Fixed effects from the linear mixed-effects model examining the effects of stimulation condition and trial on reaction time.

| Predictor | *b* | *SE* | *t* | 95% CI | *p* |
| --- | --- | --- | --- | --- | --- |
| PA | -0.036 | 0.023 | -1.55 | [-0.082, 0.010] | .120 |
| PT | -0.033 | 0.023 | -1.42 | [-0.078, 0.013] | .157 |
| FPA | 0.004 | 0.023 | 0.19 | [-0.041, 0.050] | .848 |
| FPT | -0.008 | 0.024 | -0.34 | [-0.054, 0.038] | .730 |
| Trial | 0.00053 | 0.00015 | 3.57 | [0.00024, 0.00082] | < .001 |
| PA × Trial | -0.00005 | 0.00021 | -0.22 | [-0.00046, 0.00037] | .828 |
| PT × Trial | 0.00016 | 0.00021 | 0.76 | [-0.00025, 0.00057] | .448 |
| FPA × Trial | -0.00001 | 0.00021 | -0.07 | [-0.00043, 0.00040] | .945 |
| FPT × Trial | 0.00008 | 0.00021 | 0.36 | [-0.00034, 0.00049] | .720 |

**Note.** *b* = unstandardized regression coefficient; *SE* = standard error; *CI* = confidence interval. Reaction time (RT) was measured in seconds. The control group served as the reference category. PA = Posterior alpha; PT = Posterior theta; FPA = fronto-parietal alpha; FPT = fronto-parietal theta. The model included random intercepts and random slopes for Trial for each participant.

**Supplementary Table 2.** Linear mixed-effects model predicting Discontinuity of Mind (DIM) scores from stimulation protocol, time, and their interaction.

| Predictor | *b* | *SE* | *t* | 95% CI | *p* |
| --- | --- | --- | --- | --- | --- |
| Post | 0.100 | 0.738 | 0.14 | [-1.353, 1.553] | .892 |
| PA stimulation × Post | 0.456 | 1.073 | 0.42 | [-1.655, 2.566] | .671 |
| PT stimulation × Post | -0.373 | 1.020 | -0.37 | [-2.380, 1.635] | .715 |
| FPA stimulation<br>× Post | −0.225 | 1.028 | −0.22 | [−2.247, 1.797] | .827 |
| FPT stimulation<br>× Post | −0.721 | 1.053 | −0.68 | [−2.793, 1.352] | .494 |

**Supplementary Table 3.** Linear mixed-effects model predicting Theory of Mind (TIM) scores from stimulation protocol, time, and their interaction.

| Predictor | *b* | *SE* | *t* | 95% CI | *p* |
| --- | --- | --- | --- | --- | --- |
| Post | 0.367 | 0.502 | 0.73 | [−0.621, 1.354] | .465 |
| PA stimulation × Post | −0.404 | 0.729 | −0.55 | [−1.838, 1.031] | .580 |
| PT stimulation × Post | −0.518 | 0.693 | −0.75 | [−1.883, 0.846] | .455 |
| FPA stimulation × Post | 0.290 | 0.698 | 0.41 | [−1.085, 1.664] | .679 |
| FPT stimulation × Post | −0.987 | 0.716 | −1.38 | [−2.396, 0.421] | .169 |

**Supplementary Table 4.** Linear mixed-effects model predicting Self scores from stimulation protocol, time, and their interaction.

| Predictor | *b* | *SE* | *t* | 95% CI | *p* |
| --- | --- | --- | --- | --- | --- |
| Post | −0.567 | 0.525 | −1.08 | [−1.599, 0.466] | .281 |
| PA stimulation × Post | 1.270 | 0.762 | 1.67 | [−0.230, 2.771] | .097 |
| PT stimulation × Post | 0.324 | 0.725 | 0.45 | [−1.103, 1.751] | .655 |
| FPA stimulation × Post | 0.535 | 0.730 | 0.73 | [−0.902, 1.973] | .464 |
| FPT stimulation × Post | 0.049 | 0.748 | 0.07 | [−1.424, 1.522] | .947 |

**Supplementary Table 5.**
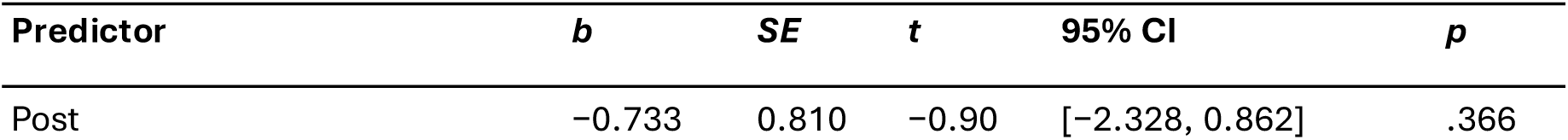
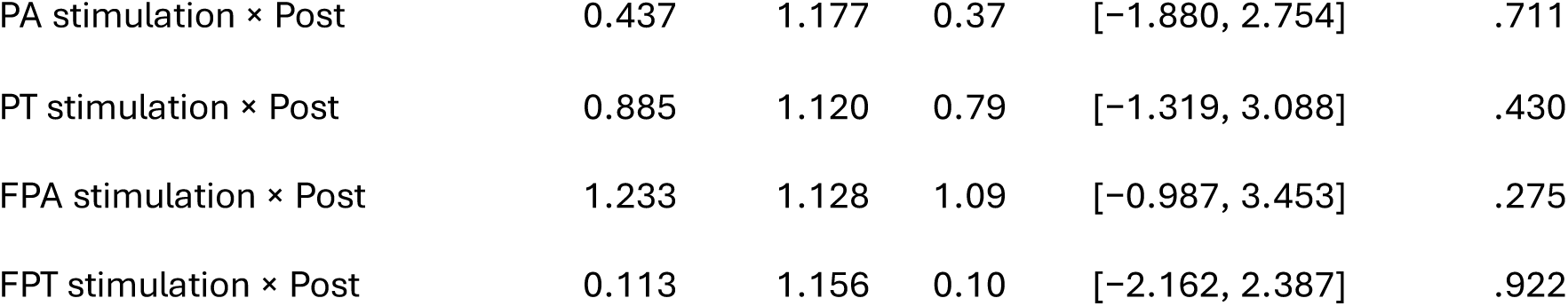
Linear mixed-effects model predicting Planning scores from stimulation protocol, time, and their interaction.

| Predictor | *b* | *SE* | *t* | 95% CI | *p* |
| --- | --- | --- | --- | --- | --- |
| Post | −0.733 | 0.810 | −0.90 | [−2.328, 0.862] | .366 |
| PA stimulation × Post | 0.437 | 1.177 | 0.37 | [−1.880, 2.754] | .711 |
| PT stimulation × Post | 0.885 | 1.120 | 0.79 | [−1.319, 3.088] | .430 |
| FPA stimulation × Post | 1.233 | 1.128 | 1.09 | [−0.987, 3.453] | .275 |
| FPT stimulation × Post | 0.113 | 1.156 | 0.10 | [−2.162, 2.387] | .922 |

**Supplementary Table 6.**
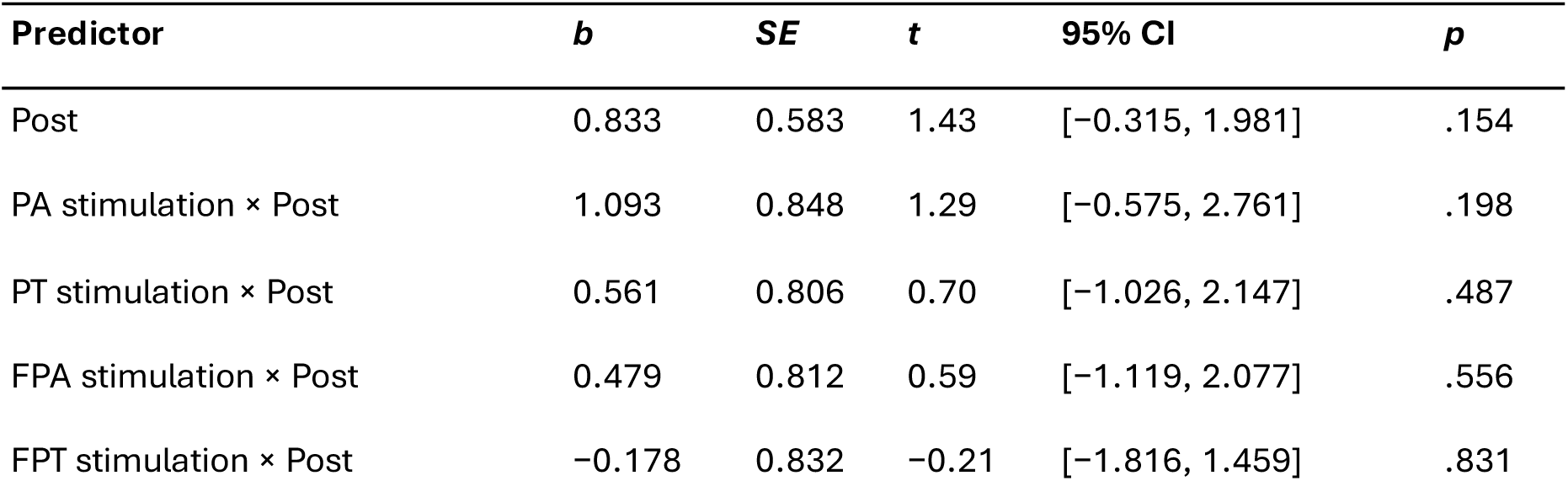
Linear mixed-effects model predicting Sleepiness scores from stimulation protocol, time, and their interaction.

| Predictor | *b* | *SE* | *t* | 95% CI | *p* |
| --- | --- | --- | --- | --- | --- |
| Post | 0.833 | 0.583 | 1.43 | [−0.315, 1.981] | .154 |
| PA stimulation × Post | 1.093 | 0.848 | 1.29 | [−0.575, 2.761] | .198 |
| PT stimulation × Post | 0.561 | 0.806 | 0.70 | [−1.026, 2.147] | .487 |
| FPA stimulation × Post | 0.479 | 0.812 | 0.59 | [−1.119, 2.077] | .556 |
| FPT stimulation × Post | −0.178 | 0.832 | −0.21 | [−1.816, 1.459] | .831 |

**Supplementary Table 7.** Linear mixed-effects model predicting Comfort scores from stimulation protocol, time, and their interaction.

| Predictor | *b* | *SE* | *t* | 95% CI | *p* |
| --- | --- | --- | --- | --- | --- |
| Post | 0.300 | 0.440 | 0.68 | [−0.566, 1.166] | .496 |
| PA stimulation × Post | −0.893 | 0.639 | −1.40 | [−2.151, 0.366] | .164 |
| PT stimulation × Post | −0.694 | 0.608 | −1.14 | [−1.891, 0.503] | .255 |
| FPA stimulation × Post | −0.956 | 0.612 | −1.56 | [−2.162, 0.249] | .120 |
| FPT stimulation × Post | −0.852 | 0.628 | −1.36 | [−2.087, 0.384] | .176 |

**Supplementary Table 8.** Linear mixed-effects model predicting Somatic Awareness (SA) scores from stimulation protocol, time, and their interaction.

| Predictor | *b* | *SE* | *t* | 95% CI | *p* |
| --- | --- | --- | --- | --- | --- |
| Intercept | 11.467 | 0.670 | 17.12 | [10.149, 12.785] | < .001 |
| PA stimulation | -0.430 | 0.973 | -0.44 | [-2.345, 1.485] | .659 |
| PT stimulation | -1.618 | 0.925 | -1.75 | [-3.439, 0.203] | .081 |
| FPA stimulation | 0.382 | 0.925 | 0.41 | [-1.439, 2.203] | .680 |
| FPT stimulation | -0.674 | 0.955 | -0.71 | [-2.553, 1.206] | .481 |
| Post | -1.233 | 0.597 | -2.06 | [-2.409, -0.057] | .040 |
| PA stimulation × Post | 1.382 | 0.868 | 1.59 | [-0.327, 3.090] | .113 |
| PT stimulation × Post | 0.294 | 0.826 | 0.36 | [-1.331, 1.919] | .722 |
| FPA stimulation × Post | 0.385 | 0.826 | 0.47 | [-1.240, 2.010] | .641 |
| FPT stimulation × Post | 0.613 | 0.852 | 0.72 | [-1.065, 2.290] | .473 |

**Note Tables 2-8.** *b* = unstandardized regression coefficient; *SE* = standard error; *CI* = confidence interval. Control group (no stimulation) and Pre stimulation session served as the reference categories. PA = Posterior alpha (Oz–Cz alpha); PT = Posterior theta (Oz–Cz theta); FPA = fronto-parietal alpha (Fz–Pz alpha); FPT = fronto-parietal theta (Fz–Pz theta).

**Supplementary Table 9.** Linear mixed-effects model predicting alpha power for electrode Fp1.

| Predictor | *b* | *SE* | *t* | 95% CI | *p* |
| --- | --- | --- | --- | --- | --- |
| Post | -0.028 | 0.111 | -0.25 | [-0.246, 0.190] | .801 |
| FPA stimulation × Post | 0.150 | 0.160 | 0.94 | [-0.166, 0.466] | .351 |
| FPT stimulation × Post | 0.285 | 0.163 | 1.74 | [-0.037, 0.607] | .082 |
| PT stimulation × Post | 0.163 | 0.164 | 1.00 | [-0.159, 0.485] | .320 |
| PA stimulation × Post | 0.484 | 0.159 | 3.05 | [0.172, 0.797] | **.002** |

**Supplementary Table 10.** Linear mixed-effects model predicting alpha power for electrode Fp2.

| Predictor | *b* | *SE* | *t* | 95% CI | *p* |
| --- | --- | --- | --- | --- | --- |
| Post | -0.005 | 0.113 | -0.05 | [-0.229, 0.218] | .963 |

| **Predictor** | ***b*** | ***SE*** | ***t*** | **95% CI** | ***p*** |
| --- | --- | --- | --- | --- | --- |
| FPA stimulation × Post | 0.136 | 0.165 | 0.82 | [−0.189, 0.462] | .410 |
| FPT stimulation × Post | 0.286 | 0.166 | 1.73 | [−0.040, 0.613] | .086 |
| PT stimulation × Post | 0.126 | 0.163 | 0.77 | [−0.196, 0.447] | .442 |
| PA stimulation × Post | 0.399 | 0.158 | 2.53 | [0.088, 0.711] | **.012** |

**Supplementary Table 11.** Linear mixed-effects model predicting alpha power for electrode F7.

| **Predictor** | ***b*** | ***SE*** | ***t*** | **95% CI** | ***p*** |
| --- | --- | --- | --- | --- | --- |
| Post | −0.044 | 0.108 | −0.40 | [−0.257, 0.169] | .686 |
| FPA stimulation × Post | 0.182 | 0.153 | 1.19 | [−0.120, 0.484] | .236 |
| FPT stimulation × Post | 0.322 | 0.156 | 2.06 | [0.014, 0.629] | **.040** |
| PT stimulation × Post | 0.233 | 0.155 | 1.50 | [−0.072, 0.537] | .134 |
| PA stimulation × Post | 0.342 | 0.150 | 2.27 | [0.046, 0.638] | **.024** |

**Supplementary Table 12.** Linear mixed-effects model predicting alpha power for electrode F3.

| **Predictor** | ***b*** | ***SE*** | ***t*** | **95% CI** | ***p*** |
| --- | --- | --- | --- | --- | --- |
| Post | −0.005 | 0.109 | −0.05 | [−0.221, 0.210] | .963 |
| FPA stimulation × Post | 0.118 | 0.154 | 0.77 | [−0.185, 0.420] | .444 |
| FPT stimulation × Post | 0.136 | 0.154 | 0.88 | [−0.168, 0.440] | .379 |
| PT stimulation × Post | 0.243 | 0.157 | 1.55 | [−0.066, 0.552] | .122 |
| PA stimulation × Post | 0.368 | 0.148 | 2.48 | [0.076, 0.660] | **.014** |

**Supplementary Table 13.** Linear mixed-effects model predicting alpha power for electrode Fz.

| **Predictor** | ***b*** | ***SE*** | ***t*** | **95% CI** | ***p*** |
| --- | --- | --- | --- | --- | --- |
| Post | −0.004 | 0.183 | −0.02 | [−0.365, 0.357] | .983 |
| FPA stimulation × Post | 0.141 | 0.262 | 0.54 | [-0.374, 0.657] | .589 |
| FPT stimulation × Post | 0.208 | 0.266 | 0.78 | [-0.316, 0.732] | .435 |
| PT stimulation × Post | 0.314 | 0.253 | 1.24 | [-0.185, 0.813] | .216 |
| PA stimulation × Post | 0.650 | 0.249 | 2.61 | [0.159, 1.140] | **.010** |

**Supplementary Table 14.** Linear mixed-effects model predicting alpha power for electrode F4.

| **Predictor** | ***b*** | ***SE*** | ***t*** | **95% CI** | ***p*** |
| --- | --- | --- | --- | --- | --- |
| Post | 0.012 | 0.101 | 0.12 | [-0.186, 0.210] | .908 |
| FPA stimulation × Post | 0.094 | 0.151 | 0.62 | [-0.203, 0.391] | .534 |
| FPT stimulation × Post | 0.165 | 0.147 | 1.12 | [-0.125, 0.454] | .264 |
| PT stimulation × Post | 0.155 | 0.145 | 1.07 | [-0.131, 0.440] | .288 |
| PA stimulation × Post | 0.464 | 0.140 | 3.31 | [0.187, 0.740] | **.001** |

**Supplementary Table 15.** Linear mixed-effects model predicting alpha power for electrode F8.

| **Predictor** | ***b*** | ***SE*** | ***t*** | **95% CI** | ***p*** |
| --- | --- | --- | --- | --- | --- |
| Post | 0.052 | 0.077 | 0.68 | [-0.099, 0.204] | .497 |
| FPA stimulation × Post | 0.007 | 0.112 | 0.06 | [-0.213, 0.226] | .952 |
| FPT stimulation × Post | 0.134 | 0.111 | 1.20 | [-0.085, 0.353] | .230 |
| PT stimulation × Post | 0.085 | 0.111 | 0.76 | [-0.134, 0.303] | .447 |
| PA stimulation × Post | 0.233 | 0.109 | 2.14 | [0.018, 0.447] | **.034** |

**Supplementary Table 16.** Linear mixed-effects model predicting alpha power for electrode T7.

| **Predictor** | ***b*** | ***SE*** | ***t*** | **95% CI** | ***p*** |
| --- | --- | --- | --- | --- | --- |
| Post | 0.102 | 0.069 | 1.48 | [-0.034, 0.238] | .141 |
| FPA stimulation × Post | −0.120 | 0.100 | −1.20 | [−0.316, 0.077] | .231 |
| FPT stimulation × Post | 0.063 | 0.099 | 0.64 | [−0.132, 0.258] | .525 |
| PT stimulation × Post | 0.019 | 0.102 | 0.19 | [−0.182, 0.219] | .853 |
| PA stimulation × Post | 0.054 | 0.098 | 0.55 | [−0.139, 0.247] | .584 |

**Supplementary Table 17.** Linear mixed-effects model predicting alpha power for electrode C3.

| **Predictor** | ***b*** | ***SE*** | ***t*** | **95% CI** | ***p*** |
| --- | --- | --- | --- | --- | --- |
| Post | −0.079 | 0.099 | −0.80 | [−0.274, 0.115] | .422 |
| FPA stimulation × Post | 0.093 | 0.145 | 0.64 | [−0.193, 0.378] | .524 |
| FPT stimulation × Post | 0.289 | 0.142 | 2.03 | [0.008, 0.569] | **.044** |
| PT stimulation × Post | 0.221 | 0.142 | 1.55 | [−0.059, 0.501] | .121 |
| PA stimulation × Post | 0.259 | 0.139 | 1.87 | [−0.014, 0.532] | .063 |

**Supplementary Table 18.**
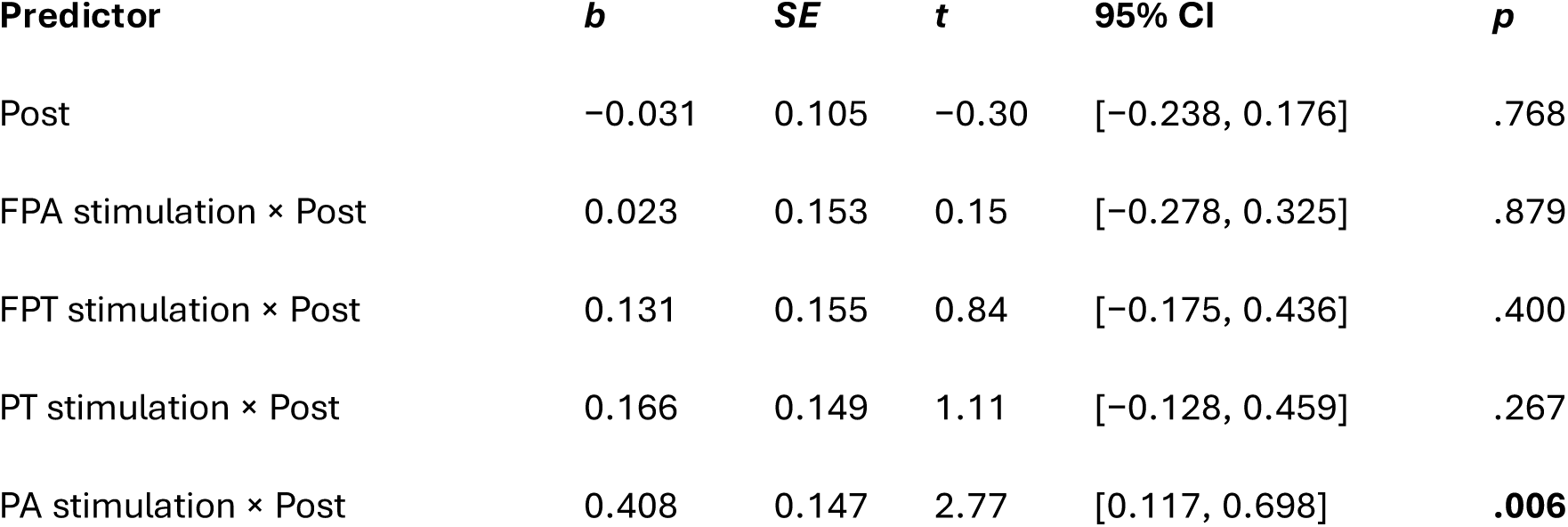
Linear mixed-effects model predicting alpha power for electrode Cz.

**Supplementary Table 19.**
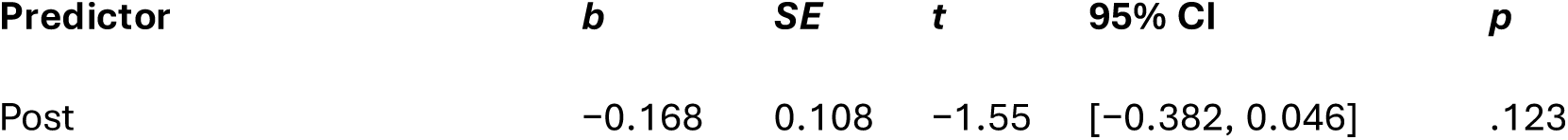
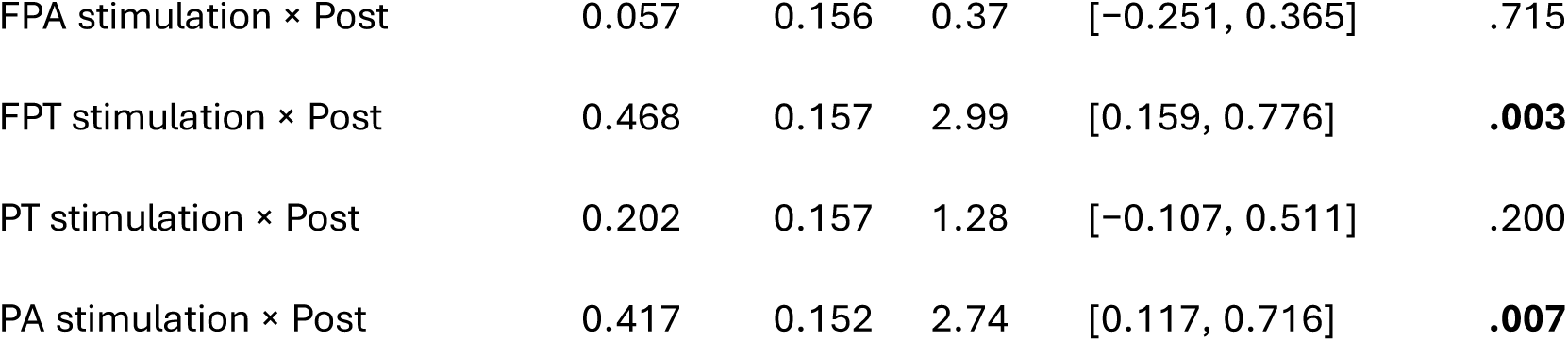
Linear mixed-effects model predicting alpha power for electrode C4.

**Supplementary Table 20.** Linear mixed-effects model predicting alpha power for electrode T8.

| **Predictor** | ***b*** | ***SE*** | ***t*** | **95% CI** | ***p*** |
| --- | --- | --- | --- | --- | --- |
| Post | 0.046 | 0.079 | 0.58 | [−0.110, 0.202] | .566 |
| FPA stimulation × Post | −0.009 | 0.116 | −0.08 | [−0.237, 0.220] | .940 |
| FPT stimulation × Post | 0.125 | 0.115 | 1.09 | [−0.100, 0.351] | .276 |
| PT stimulation × Post | 0.148 | 0.113 | 1.31 | [−0.075, 0.371] | .192 |
| PA stimulation × Post | 0.191 | 0.111 | 1.71 | [−0.028, 0.410] | .088 |

**Supplementary Table 21.** Linear mixed-effects model predicting alpha power for electrode P7.

| **Predictor** | ***b*** | ***SE*** | ***t*** | **95% CI** | ***p*** |
| --- | --- | --- | --- | --- | --- |
| Post | 0.217 | 0.163 | 1.33 | [−0.104, 0.537] | .185 |
| FPA stimulation × Post | −0.186 | 0.239 | −0.78 | [−0.656, 0.285] | .438 |
| FPT stimulation × Post | −0.028 | 0.233 | −0.12 | [−0.487, 0.431] | .904 |
| PT stimulation × Post | 0.134 | 0.239 | 0.56 | [−0.336, 0.605] | .575 |
| PA stimulation × Post | 0.224 | 0.231 | 0.97 | [−0.230, 0.678] | .333 |

**Supplementary Table 22.** Linear mixed-effects model predicting alpha power for electrode P3.

| **Predictor** | ***b*** | ***SE*** | ***t*** | **95% CI** | ***p*** |
| --- | --- | --- | --- | --- | --- |
| Post | 0.151 | 0.188 | 0.80 | [−0.219, 0.522] | .422 |
| FPA stimulation × Post | −0.080 | 0.268 | −0.30 | [−0.609, 0.448] | .765 |
| FPT stimulation × Post | 0.250 | 0.274 | 0.91 | [−0.289, 0.790] | .362 |
| PT stimulation × Post | 0.110 | 0.269 | 0.41 | [−0.420, 0.639] | .684 |
| PA stimulation × Post | 0.291 | 0.260 | 1.12 | [−0.221, 0.804] | .264 |

**Supplementary Table 23.** Linear mixed-effects model predicting alpha power for electrode Pz.

| **Predictor** | ***b*** | ***SE*** | ***t*** | **95% CI** | ***p*** |
| --- | --- | --- | --- | --- | --- |
| Post | −0.066 | 0.150 | −0.44 | [−0.362, 0.230] | .661 |
| FPA stimulation × Post | 0.311 | 0.211 | 1.47 | [−0.104, 0.726] | .142 |
| FPT stimulation × Post | 0.372 | 0.215 | 1.73 | [−0.050, 0.795] | .084 |
| PT stimulation × Post | 0.087 | 0.215 | 0.41 | [−0.335, 0.510] | .684 |
| PA stimulation × Post | 0.601 | 0.209 | 2.87 | [0.189, 1.012] | **.004** |

**Supplementary Table 24.**
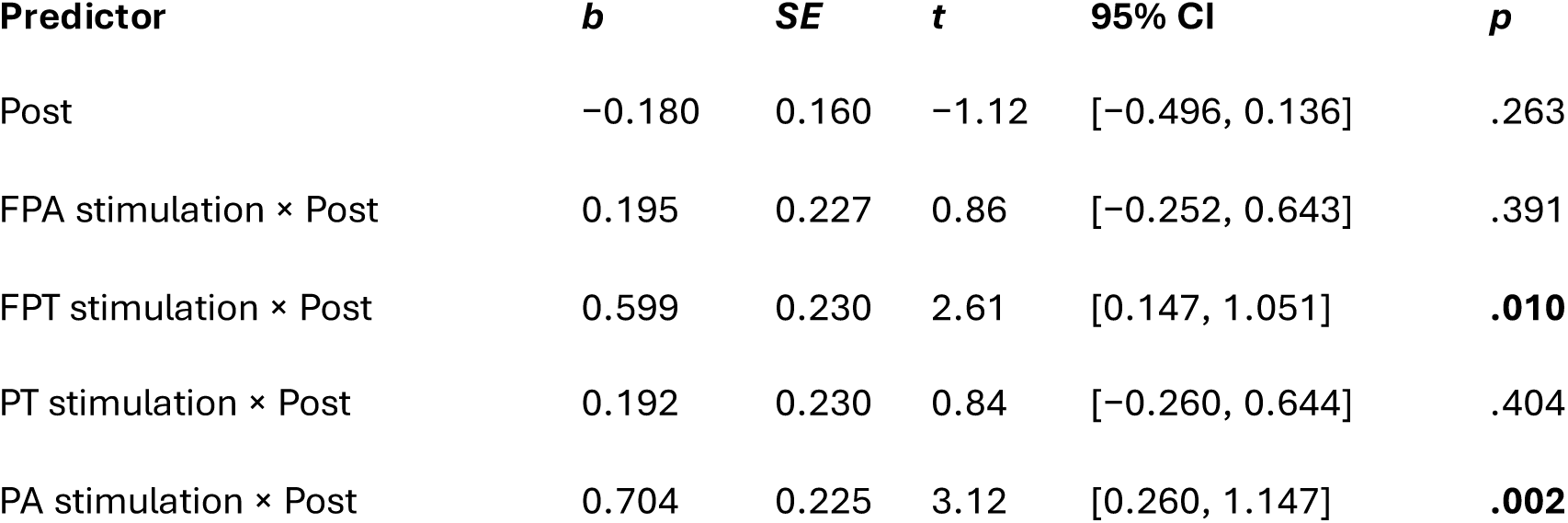
Linear mixed-effects model predicting alpha power for electrode P4.

| **Predictor** | ***b*** | ***SE*** | ***t*** | **95% CI** | ***p*** |
| --- | --- | --- | --- | --- | --- |
| Post | −0.180 | 0.160 | −1.12 | [−0.496, 0.136] | .263 |
| FPA stimulation × Post | 0.195 | 0.227 | 0.86 | [−0.252, 0.643] | .391 |
| FPT stimulation × Post | 0.599 | 0.230 | 2.61 | [0.147, 1.051] | **.010** |
| PT stimulation × Post | 0.192 | 0.230 | 0.84 | [−0.260, 0.644] | .404 |
| PA stimulation × Post | 0.704 | 0.225 | 3.12 | [0.260, 1.147] | **.002** |

**Supplementary Table 25.** Linear mixed-effects model predicting alpha power for electrode O1.

| **Predictor** | ***b*** | ***SE*** | ***t*** | **95% CI** | ***p*** |
| --- | --- | --- | --- | --- | --- |
| Post | 0.052 | 0.322 | 0.16 | [−0.582, 0.687] | .871 |
| FPA stimulation × Post | 0.088 | 0.475 | 0.19 | [-0.847, 1.022] | .853 |
| FPT stimulation × Post | 0.366 | 0.466 | 0.78 | [-0.552, 1.284] | .433 |
| PT stimulation × Post | 0.316 | 0.461 | 0.69 | [-0.591, 1.223] | .494 |
| PA stimulation × Post | 1.353 | 0.453 | 2.99 | [0.462, 2.244] | **.003** |

**Supplementary Table 26.** Linear mixed-effects model predicting alpha power for electrode Oz.

| **Predictor** | ***b*** | ***SE*** | ***t*** | **95% CI** | ***p*** |
| --- | --- | --- | --- | --- | --- |
| Post | -0.087 | 0.279 | -0.31 | [-0.635, 0.462] | .756 |
| FPA stimulation × Post | 0.218 | 0.402 | 0.54 | [-0.574, 1.009] | .589 |
| FPT stimulation × Post | 0.378 | 0.400 | 0.95 | [-0.410, 1.167] | .345 |
| PT stimulation × Post | 0.675 | 0.393 | 1.72 | [-0.098, 1.448] | .087 |
| PA stimulation × Post | 1.252 | 0.383 | 3.27 | [0.498, 2.007] | **.001** |

**Supplementary Table 27.**
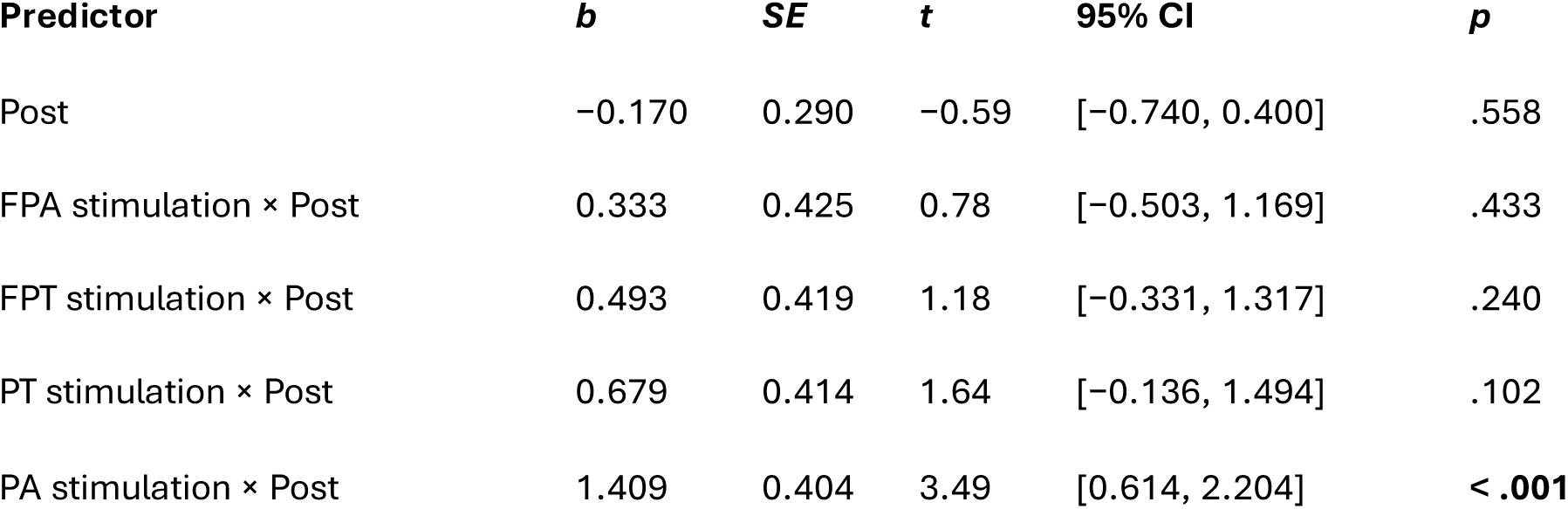
Linear mixed-effects model predicting alpha power for electrode O2.

| **Predictor** | ***b*** | ***SE*** | ***t*** | **95% CI** | ***p*** |
| --- | --- | --- | --- | --- | --- |
| Post | -0.170 | 0.290 | -0.59 | [-0.740, 0.400] | .558 |
| FPA stimulation × Post | 0.333 | 0.425 | 0.78 | [-0.503, 1.169] | .433 |
| FPT stimulation × Post | 0.493 | 0.419 | 1.18 | [-0.331, 1.317] | .240 |
| PT stimulation × Post | 0.679 | 0.414 | 1.64 | [-0.136, 1.494] | .102 |
| PA stimulation × Post | 1.409 | 0.404 | 3.49 | [0.614, 2.204] | **< .001** |

**Supplementary Table 28.** Linear mixed-effects model predicting alpha power for electrode P8.

| **Predictor** | ***b*** | ***SE*** | ***t*** | **95% CI** | ***p*** |
| --- | --- | --- | --- | --- | --- |
| Post | -0.031 | 0.170 | -0.18 | [-0.365, 0.304] | .857 |
| FPA stimulation × Post | −0.039 | 0.239 | −0.16 | [−0.510, 0.432] | .870 |
| FPT stimulation × Post | 0.228 | 0.243 | 0.94 | [−0.251, 0.707] | .349 |
| PT stimulation × Post | 0.433 | 0.241 | 1.80 | [−0.042, 0.907] | .074 |
| PA stimulation × Post | 0.621 | 0.235 | 2.64 | [0.158, 1.084] | **.009** |

**Note.** Tables 9–28 report the fixed effects of **Time** and the **Group × Time** interaction from separate linear mixed-effects models fitted for each EEG electrode. *b* = unstandardized regression coefficient; *SE* = standard error; CI = confidence interval. The Control group (no stimulation) and the Pre stimulation session served as the reference categories. PA = Posterior alpha (Oz–Cz alpha); PT = Posterior theta (Oz–Cz theta); FPA = fronto-parietal alpha (Fz–Pz alpha); FPT = fronto-parietal theta (Fz–Pz theta).

**Supplementary Table 29.** FDR-corrected p-values for linear mixed-effects model predicting alpha power (Stimulation x Post Interaction)

| **Electrode** | **FPA × Post** | **FPT × Post** | **PT × Post** | **PA × Post** |
| --- | --- | --- | --- | --- |
| Fp1 | .952 | .484 | .446 | **.017** |
| Fp2 | .952 | .484 | .481 | .103 |
| F7 | .952 | .484 | .596 | **.045** |
| F3 | .906 | .484 | .523 | **.008** |
| Fz | .952 | .062 | .481 | **.015** |
| F4 | .906 | .096 | .596 | **.010** |
| F8 | .906 | .245 | .596 | **.020** |
| T7 | .906 | .245 | .533 | **.010** |
| C3 | .906 | .484 | .481 | **.018** |
| Cz | .952 | .484 | .523 | **.015** |
| C4 | .952 | .484 | .617 | **.010** |
| T8 | .906 | .484 | .446 | **.008** |
| P7 | .906 | .484 | .446 | **.008** |
| P3 | .906 | .245 | .720 | **.013** |

| Electrode | FPA × Post | FPT × Post | PT × Post | PA × Post |
| --- | --- | --- | --- | --- |
| Pz | .952 | .484 | .720 | .294 |
| P4 | .906 | .218 | .446 | .078 |
| P8 | .906 | .484 | .446 | **.021** |
| O1 | .906 | .218 | .446 | **.034** |
| Oz | .906 | .552 | .853 | .583 |
| O2 | .906 | .904 | .676 | .350 |

**Note.** Values are false discovery rate (FDR)-corrected $p$ -values for the Group × Post interaction terms. Significant effects ( $p < .05$ ) are shown in bold.

